# Sex-Specific Genetic Architecture Of Behavioral Traits In A Spider

**DOI:** 10.1101/2020.07.28.224675

**Authors:** Simona Kralj-Fišer, Jutta M. Schneider, M. Kuntner, Francisco Garcia-Gonzalez

## Abstract

Sex differences in behavioral traits are common, but we know little about the role of sexual selection in shaping these traits. Estimating sex-specific genetic effects and cross-sex genetic correlations can provide insights into sex-specific selection and on whether evolution can shape independent expression of behavioral traits across the sexes. We conducted a quantitative genetic study in a sexually-size-dimorphic spider, *Larinioides sclopetarius*, which exhibits sex differences in adult life-styles. We observed pedigreed spiders for aggression, activity, exploration and boldness, and used animal models to disentangle genetic and environmental influences on these behaviors. We detected higher additive genetic variances in activity and aggression in males compared to females, but no sex differences in quantitative genetic estimates for exploration and boldness. The estimated mean cross-sex genetic correlation in all traits were close to zero suggesting these traits could have flexibility for sex-independent evolution. We note however, that the 95% credible intervals of cross-sex genetic correlation are large, and thus estimates uncertain. Our results imply that individual variation in aggression and activity might stem from sex-specific selection acting on these traits. The estimates of sex-specific additive genetic variation and cross-sex genetic correlation suggests their further sex-independent evolution. Taken together, our results support the notion that sexual selection can play an important role shaping behavioral traits.

## INTRODUCTION

Individuals commonly differ in behaviors that are consistent over time and/or context, i.e. personality traits, coping styles or temperament (Gosling 2001; Koolhaas et al. 1999; Réale et al. 2007). Theoretically, consistent variation in behavior has been explained by spatio-temporal variation in selective pressures often generated by state-dependent positive feedback loops or negative frequency dependent selection (Wolf et al. 2007; Wolf and Weissing 2010; Sih et al. 2015). However, one major source of variation in selective pressures can be sex-specific selection (Schuett et al. 2010). Given that behaviors can act as both targets and mediators of sexual selection processes, it is important to understand sex specificity in the genetic architectures of behavioral traits.

Behavioral traits can affect an individual’s survival and reproductive success (Biro and Stamps 2008), and the behavioral strategies to maximize fitness may strongly differ between sexes (Fairbairn et al. 2007; Schuett et al. 2010). Males and females commonly differ in the mean levels of behavior, e.g. males are on average more aggressive and bolder than females (Schuett et al. 2010; e.g. Kralj-Fišer et al. 2017; Kaiser et al. 2018). Sexes may also differ in the degree of behavioral repeatability, i.e. a ratio between among- and within-individual variance in a trait, with males in general exhibiting higher repeatability in their behavior than females (Nakagawa et al. 2007; Bell et al. 2009). Schuett et al. (2010) suggested that repeatability signals predictability, which might have been selected for through mate choice and male-male competition. Despite emerging links between aspects of individual behavioral variation (mean levels, repeatability) and sexual selection (mate choice, intra-sexual competition), the role of sexual selection in the evolution of individual behavioral variation has been rarely explored empirically, though it has recently gathered increased attention (Hämäläinen et al. 2018; Immonen et al. 2018; Tarka et al. 2018). A critical first step to understand the potential role of sex-specific selection on the evolution of consistent individual behavioral differences is to identify the heritability and underlying genetic architectures (sex-specific genetic variances, genetic covariance between sexes) of behavioural traits, along with the determination of whether behavioral expression differs between the sexes.

Females and males share the same genes apart from those on heteromorphic sex chromosomes. Thus, the sexes share the genetic basis for most homologous traits. If selection pressures on those traits are opposing in males versus females, shared genetic variance may constrain one or both sexes from reaching their phenotypic optima, setting the stage for intra-locus sexual conflict. Theoretically, at least a partial resolution to intra-locus sexual conflict is required for the sex-independent trait expression and evolution of sexual dimorphism. To quantify the amount of sex-specific genetic variance and genetic divergence of the sexes, the cross-sex genetic correlation between homologous male and female traits is commonly estimated as 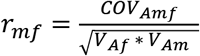, where *COV_Amf_* is the additive genetic covariance between the sexes, and *V_Am_* and *V_Af_* are additive genetic variances of males and females, respectively (Lande 1980). The cross-sex genetic correlation measures the extent of similarity between the additive alleles when expressed in both sexes (Lande 1980; Bonduriansky and Chenoweth 2009). When *r_mf_* is close to one, a shared trait is assumed to be controlled by a common genetic architecture in both sexes. On the other hand, when *r_mf_* aproaches zero, the two sexes differ in the genetic architecture for the shared trait, or differ in allele expression. Sex-independent evolution in a trait is possible through low additive genetic covariance between the sexes, sex differences in additive genetic variances, or both (Lande 1980; Lynch and Walsh 1998).

Most studies show that behavioral traits are moderately heritable, though heritability estimates often differ between traits and taxonomic groups (reviewed in van Oers and Sinn 2013; Dochtermann et al. 2019). Several studies have demonstrated among-population variation in heritability estimates of individual behavioral traits (Bell, 2005; Dingemanse et al. 2009), but so far, few have investigated within-population, and specifically, between-sex variation in genetic variances (but see Han and Dingemanse 2017; White et al. 2019). For example, in the southern field crickets, *Gryllus bimaculatus*, where males are more aggressive and more explorative than females, males have higher genetic variance for both behaviors, and there is a low cross-sex genetic correlation for aggression implying a degree of genetic independence for these traits between males and females, and a resolution of sexual conflict to some extent (Han and Dingemanse 2017). Furthermore, a recent study in an orb-web spider, *Nuctenea umbratica*, found higher heritability of aggression in males compared to females, though females and males did not differ in additive genetic variation of this trait (Kralj-Fišer et al. 2019). The two sexes, however, showed no differentiation in genetic architecture of activity (Kralj-Fišer et al. 2019). Similarly, the quantitative genetic study by White and colleagues 2019 suggested that behavioral traits related to risk-taking in the Trinidadian guppy, *Poecilia reticulata*, where males are bolder, lack sex-specific genetic architecture within traits. Namely, the sexes did not differ in the amount of sex-specific genetic variances and cross-sex genetic correlations within traits did not differ from unity (White et al. 2019). Nonetheless, there was a weak evidence of sex-specificity in genetic correlations between traits (White et al. 2019). Additional studies are needed, however, to gain a more general insight into the sex-specificity of behavioral genetic architectures, sex-specific selection and the sexual conflict dynamics of individual behavioral variation.

In this study we examine sex-specific heritability estimates of a series of behavioral traits, namely aggression, boldness, activity and exploration in a novel environment in a spider model, *Larinioides sclopetarius*. We used data generated by Kralj-Fišer and Schneider’s (2012) study, in which the above behaviors were measured in males and females. In that study, the heritability of the different behaviors was assessed using parent-offspring regression, but combined across both sexes (Kralj-Fišer and Schneider 2012). Here, we test whether females and males differ in quantitative genetic estimates in these traits. We partition the observed phenotypic variance (*V_P_*) in each trait into additive genetic (*V_A_*), common environment/maternal (*V_CE/M_*), permanent environment (*V_PE_*) and residual variances (*V_R_*). In addition, we calculated two mean-standardized evolvability measures, the coefficient of additive genetic variation and its square, as well as the coefficient of common environment/maternal effect variation, permanent environment effect variation and the coefficient of residual variation. These values enable the comparison of components of phenotypic variance and evolvability among different traits, sex and taxa (Houle 1992; Hansen et al. 2011; Garcia-Gonzalez et al. 2012). Furthermore, we estimated the cross-sex genetic correlations (*r_mf_*) of the four traits.

*Larinioides* spiders are sexually size dimorphic and exhibit large sex differences in their adult life styles, suggesting sex-specific selection pressures. Females are territorial, sit-and-wait predators, whereas males cease web building after reaching maturity; adult males wander around in search of mates and feed mostly commensally in female webs. Based on these sex differences we could predict that males exhibit higher mean levels of activity and exploration, but a previous laboratory study on this species failed to support this expectation (Kralj-Fišer and Schneider 2012). However, that study also found higher mean repeatability in males compared to females in activity and exploration, but not in boldness (activity; R♂ = 0.894, R♀ = 0.401; exploration, R♂ = 0.734, R♀ = 0.530; boldness, R♂ = 0.798, R♀ = 0.824; Kralj-Fišer and Schneider 2012). Given that the repeatability of a trait represents an upper limit for its heritability (Falconer and Mackay 1996; Lynch and Walsh 1998), we expect to find that activity and exploration would have higher heritability estimates in males than in females. Furthermore, we predict that selection on activity and exploration might be stronger for males (finding mates) than for females, who would benefit more from saving energy for offspring production, as they do not need to move to feed or find mates. Thus, we would also expect sex-specificity in the genetic architectures of these traits. We would further expect that males, who move more and may thus be more conspicuous than females, have been under stronger selection for boldness and thus may potentially have different underlying genetic architecture of the trait compared to females.

*Larinioides* males commonly fight for access to mates, while female disputes are rare (Kralj-Fišer and Schneider 2012). This was observed also under laboratory conditions, where males were generally more aggressive than females (Kralj-Fišer and Schneider 2012). In males, intra-sex aggression enhances access to mates and has been likely shaped by sexual selection. In females, aggression toward same-sex conspecifics serves to defend their territory (web) and foraging patch, and overt aggression may have high fitness costs due to injuries and death (Kralj-Fišer and Schneider 2012). The results from previous laboratory experiments imply that aggressive males father more offspring than non-aggressive ones, whereas female aggression is not related to fecundity (Kralj-Fišer et al. 2013). Based on the repeatability estimates for aggression found in that study (R♂ = 0.864, R♀ = 0.772), we expected that the sexes may not greatly differ in heritability for this trait (Kralj-Fišer and Schneider 2012). Nevertheless, we predict to find some sex differences in the underlying genetic architecture, because males and females seem to differ in their phenotypic optima for aggression.

## MATERIAL AND METHODS

### Study animals

*Larinioides sclopetarius* is a nocturnal, Holarctic orb-weaving spider, commonly found in high densities on human constructions near water bodies (Heiling and Herberstein 1998). Individuals build webs adjacent to one another, but retain territorial and aggressive habits. We collected subadult *L. sclopetarius* along riverbanks and bridges in Hamburg (Germany) in September 2010, then transferred them to the laboratory to be reared to adulthood in 200 ml plastic cups and fed with ad libitum *Drosophila sp*. flies. Adult females are larger than males with size dimorphism index of 0.85 (Turk et al. 2019), therefore we accordingly adjusted the food regime upon maturation; adult females (*N*=30) that were placed into plastic frames (36 x 36 x 6 cm) were fed with *Calliphora sp*, whereas males (*N*=31) remained in the 200 ml cups under the same feeding treatment (ad libitum *Drosophila sp.*). We fed the spiders twice a week, misted their webs with water using a spray bottle five days a week, and kept them at room temperature under 10:14 (LD) conditions. Research on spiders is not restricted by the animal protection law in EU. We collected minimal number of individuals in the field to conduct the study. Natural populations of tested spiders are abundant in the field, and their populations are not at risk. Spiders were not harmed in any way.

### Experimental Design

We used the data from Kralj-Fišer and Schneider (2012), where additional details may be found. Kralj-Fišer and Schneider (2012) observed the behavior of adult male and female spiders in a series of standardized personality tests designed to test aggression, activity, exploration and boldness. Each individual from the parental generation was tested twice in each of the test situations. Aggression was tested by placing two individuals of the same sex approximately 5 cm from each other, and their aggression related behaviors, i.e. approaching, web shaking, attacking, chasing and biting, were recorded for 20 min. Opponents were mass-matched to the average difference of 15.11% and 7.93% in females and males, respectively. Activity and exploration were tested in a novel environment. A focal spider was gently placed in an unfamiliar plastic box (11 x 11 x 6 cm) using a paintbrush, and observed for 300 seconds. When positioned in the box, the spider started walking within the box. The latency to the first pause in walking after being placed into the box was recorded (activity). The pause was considered when the spider stopped moving for at least 2 seconds. Thereafter, the latency to move again after the first halt (exploration) was recorded. In both cases, 300 seconds were taken as the maximum latency. Boldness was measured as a response to a simulated predator attack. A spider was placed in the plastic container (11 x 11 x 6 cm) using a soft brush. Then, a predator attack was simulated by shuddering the container until the spider feigned death – a spider posture very similar to that of a dead spider. The time that elapsed between the start of death feigning to the first move afterwards was used as boldness. 300 seconds were taken as the maximum duration.

The animals were then mated assortatively by aggressiveness type. 29 females produced at least one egg-sac that we collected and stored in a climate chamber at 25°C until hatching. Spiderlings (from the first clutch) were kept together until their second molt (they die if separated before); thereafter they were separated into individual cups. These offspring were reared under standard conditions as described above. After reaching maturity, their behaviors were measured in the same way as in their parents, but once in each of the tests. The aim was to test 10 full-siblings: 5 males and 5 females per family (N = 29 families), but some clutches were sex-biased. 262 spiderlings (134 sons, 128 daughters) distributed among the families in the breeding design (mean ± SE of 4.963 ± 0.189 daughters, and 4.741 ± 1.074 sons, per family) were tested.

We note that in some contexts, assortative mating can potentially bias estimates of variance components and heritability. However, if the phenotypes of all individuals including parents are included in an animal model (Walsh and Lynch 2018), as is the case here, assortative mating in fact serves to increase the precision of the estimates (Michael Morrissey unpublished; see Kralj-Fišer et al. 2019).

### Analyses

Selection works simultaneously on multiple (correlated) traits rather on a single trait (Lande and Arnold 1983). Therefore, exploring sex-specific genetic (co)variance in a multivariate sense that considers the among-trait covariance structure is desirable (Walsh and Blows 2009; Wyman et al. 2013) While the framework for the multivariate approach is available, such studies are rare (e.g. Gosden et al. 2012; White et al. 2019) which may be mainly due to difficulty of recording multiple behaviors in a large number of individuals required in the quantitative genetic studies. In our case this was precisely a constraint and therefore we asses genetic architectures on a trait by trait basis, which still gives an informative and useful, though perhaps restricted, compared to a fully multivariate approach, view on trait evolution.

Behavioral traits, aggression, activity, exploration, and boldness were z-scored using overall means and standard deviations. We used animal models to assess sex-specific quantitative genetic parameters and cross-sex genetic correlation of four behaviors (Wilson et al. 2010). Animal models are mixed-effects models that decompose phenotypic variance into genetic and environmental effects (Kruuk and Hadfield 2007). We ran Markov Chain Monte Carlo Linear Mixed Models using the package MCMCglmm in R (version 3.5.1., R Core Team, 2013, Hadfield 2010) to partition the observed phenotypic variance (*V_P_*) in a trait into additive genetic (*V_A_*), common environment/maternal (*V_CE/M_*), permanent environment – container ID (*V_PE_*) accounting for the repeated measures of each individual, and residual variances (*V_R_*). We used the command *us* to enable estimation of sex-specific additive genetic variance in a trait (females’ additive genetic variance = *V_Af_*; males’ additive genetic variance = *V_Am_*) as well as an assessment of the additive genetic covariance in a trait between males and females (*COV*_*Am*f_) and thus cross-sex genetic correlations (*r_mf_*). This was calculated as 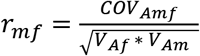 (Lande 1980). We constrained *V_PE_* and *V_R_* cross-sex covariances to zero using the command *idh*. To define genetic relatedness between individuals, we constructed a pedigree containing each individual included in the experiments. Parents of the P (parental) generation were marked as NA, as P individuals were field-collected, and their family tree was unknown. Spiderlings from the same clutch could not be separated before the second moult, thus siblings shared early common environmental effects. Sibling phenotypes, however, also shared the effects of the same mother (maternal effects). Thus, our study design did not distinguish the components of maternal from common environment effects.

We estimated the narrow sense heritability in each trait as 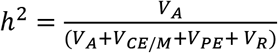 with 95% credible intervals (CIs) for females and males, separately. In the estimation of the heritability of aggressiveness we additionally included contest ID (*V*_C_) as a random effect because aggressiveness was scored simultaneously for two individuals in dyadic contests. We then estimated narrow-sense heritability in aggressiveness as 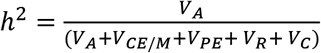.

Additionally, we calculated two mean-standardized evolvability measures, the coefficient of additive genetic variation, 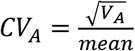, and its square, 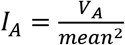, the coefficient of common environment/maternal effect variation as 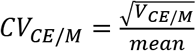, the coefficient of permanent environment effect variation as 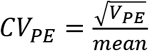 and the coefficient of residual variation as 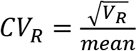 (Houle, 1992; Garcia-Gonzalez et al. 2012). *CV_A_, I_A_, CV_M/CE_, CV_PE_* and *CV_R_* and their 95% CI were calculated using the raw data (unstandardized) and phenotypic means, which were always obtained from both the parental and offspring generations. We then calculated the mean differences between the female and male in *CV_A_, I_A_, CV_M/CE_, CV_PE_* and *CV_R_* for all posterior estimates and obtained the 95% CI for these differences.

We run models with three different priors to check if they are uninformative (see in Supplementary material 1). We checked convergence and mixing properties by visual inspection of the chains and checked the autocorrelation values. We ran Heidelberger and Welch’s convergence diagnostics to verify that the number of iterations was adequate for chains to achieve convergence.

## RESULTS

### Aggression

Females exhibit lower mean aggression compared to males (post. mean difference = −0.929, 95% credible interval (CI) = [−1.217, −0.641]; p < 0.001; Figure 1). The quantitative genetic estimates for females and males are given in Tables 1 and 2. Figure 2A represents the total phenotypic variance (*V_P_*) in aggression scores decomposed into additive genetic variance (*V_A_*), common environment/ maternal effects variance (*V_CE/M_*), permanent environment variance (*V_PE_*) and residual variance (*V_R_*) in females and males separately. Both the additive genetic variation (*V_A_*) and residual variation (*V_R_*) of aggression are significantly lower in females than in males (*V_A_*, post. mean difference = −0.469, 95% CI [−0.898, −0.071]; *V_R_*, post. mean difference = – 0.568, 95% CI [−0.919, −0.241]). Also the coefficient of residual variation (C*V_R_*) is lower in females than males (C*V_R_*, post. mean difference = – 0.370, 95% CI [−0.589, −0.169]), however we found no significant sex difference in *CV_A_* (Table 2). Furthermore, variances due to maternal effect (*V_CE/M_*) and permanent effect (*V_PE_*), as well as heritability estimate (*h^2^*) are lower in females; differences, however, are not statistically significant, though based on the CIs the existing differences cannot be ruled out (*V_CE/M_*, post. mean difference = −0.127, 95% CI [−0.453, 0.039]; *V_PE_*, post. mean difference = −0.120, 95% CI [−0.439, 0.037]; *h^2^*, post. mean difference = −0.206, 95% CI [−0.476, 0.066]). Yet, we found no sex differences in coefficients of maternal effect (*CV_CE/M_*) and permanent effect (*CV_PE_*) variances (Table 2). We also found no differences between females and males in evolvability (*I_A_*) of aggression (Table 2). The mean additive genetic covariance between sexes is estimated to 0.015 with 95% CI [−0.055, 0.097]. The cross-sex genetic correlation is 0.170 with 95% CI [−0.557, 0.852].

**Figure 1.**
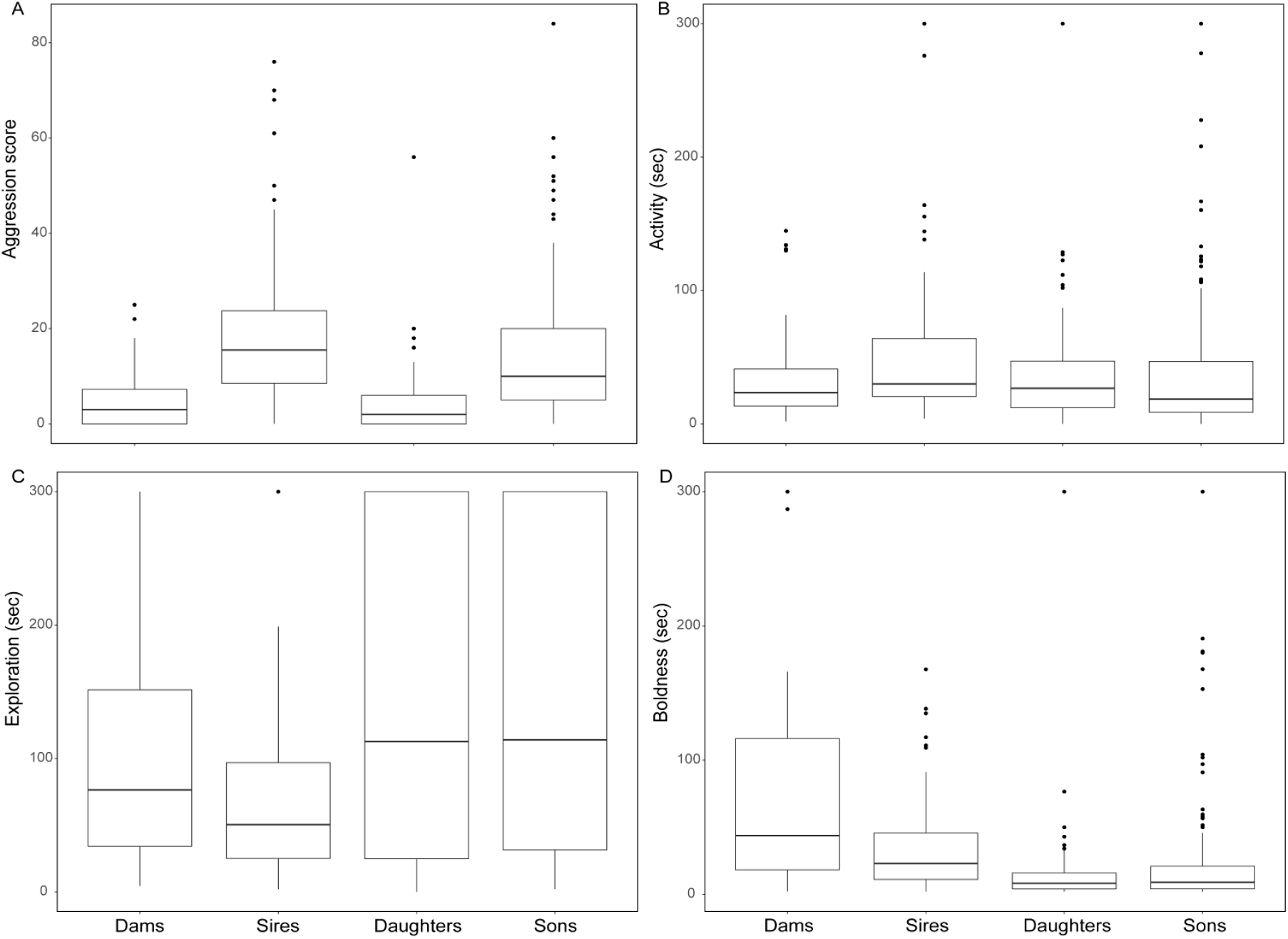
Boxplots show the median and 25^th^ and 75^th^ percentiles of behaviors in dams, sires, daughters and sons. A) aggression scores, B) activity – initial duration of movement after being introduced into a novel environment, C) exploration – the latency to move again after the first stop, and D) boldness – the time that elapsed between the start of death feigning after simulated predator attack to the first move afterwards.

**Table 1.** Estimates (posterior mean (95% credible interval) of the additive genetic variance (*V_A_*), common environment/ maternal effects variance (*V_CE/M_*), permanent environment variance (*V_PE_*), residual variance (*V_R_*), and heritability (*h^2^*). Behaviors were mean centered using overall means and standard deviations before the analyses.

| Behaviour | Sex | $V_A$<br>mean (95% CI) | $V_{CE/M}$<br>mean (95% CI) | $V_{PE}$<br>mean [95% CI] | $V_R$<br>mean [95% CI] | $h^2$<br>mean (95% CI) |
| --- | --- | --- | --- | --- | --- | --- |
| Aggression | ♀ | 0.020 [<0.001, 0.062] | 0.008 [<0.001, 0.030] | 0.008 [<0.001, 0.028] | 0.167 [0.106, 0.229] | 0.082 [<0.001, 0.246] |
|  | ♂ | 0.489 [<0.001, 0.919] | 0.135 [<0.001, 0.450] | 0.128 [<0.001, 0.433] | 0.735 [0.412, 1.076] | 0.288 [0.070, 0.506] |
|  | ♀ minus ♂ | **-0.469 [-0.898, -0.071]** | *-0.127 [-0.453, 0.039]* | *-0.120 [-0.439, 0.037]* | **-0.568 [-0.919, -0.241]** | *-0.206 [-0.476, 0.066]* |
| Activity | ♀ | 0.017 [<0.001, 0.064] | 0.079 [<0.001, 0.197] | 0.106 [<0.001, 0.287] | 0.331 [0.204, 0.455] | 0.036 [<0.001, 0.135] |
|  | ♂ | 0.507 [<0.001, 1.170] | 0.817 [0.219, 1.388] | 0.705 [0.227, 1.160] | 0.315 [0.171, 0.490] | 0.290 [<0.001, 0.611] |
|  | ♀ minus ♂ | *-0.491 [-1.185, 0.036]* | **-0.738 [-1.323, -0.130]** | **-0.599 [-1.087, -0.092]** | **0.016 [-0.206, 0.221]** | *-0.254 [-0.627, 0.063]* |
| Exploration | ♀ | 0.060 [<0.001, 0.224] | 0.138 [<0.001, 0.340] | 0.335 [<0.001, 0.620] | 0.512 [0.262, 0.818] | 0.056 [<0.001, 0.208] |
|  | ♂ | 0.223 [<0.001, 0.607] | 0.082 [<0.001, 0.271] | 0.456 [0.227, 0.783] | 0.443 [0.243, 0.689] | 0.181 [<0.001, 0.467] |
|  | ♀ minus ♂ | -0.163 [-0.635, 0.210] | 0.056 [-0.239, 0.351] | -0.121 [-0.635, 0.427] | 0.069 [-0.302, 0.466] | -0.125 [-0.487, 0.177] |
| Boldness | ♀ | 0.103 [<0.001, 0.396] | 0.129 [<0.001, 0.484] | 0.365 [<0.001, 0.625] | 0.764 [0.526, 1.028] | 0.071 [<0.001, 0.254] |
|  | ♂ | 0.049 [<0.001, 0.175] | 0.060 [<0.001, 0.182] | 0.312 [<0.001, 0.538] | 0.381 [0.188, 0.625] | 0.060 [<0.001, 0.211] |
|  | ♀ minus ♂ | 0.055 [-0.199, 0.419] | 0.069 [-0.210, 0.482] | 0.053 [-0.385, 0.473] | **0.383 [0.030, 0.734]** | 0.012 [-0.223, 0.264] |

**Table 2.**
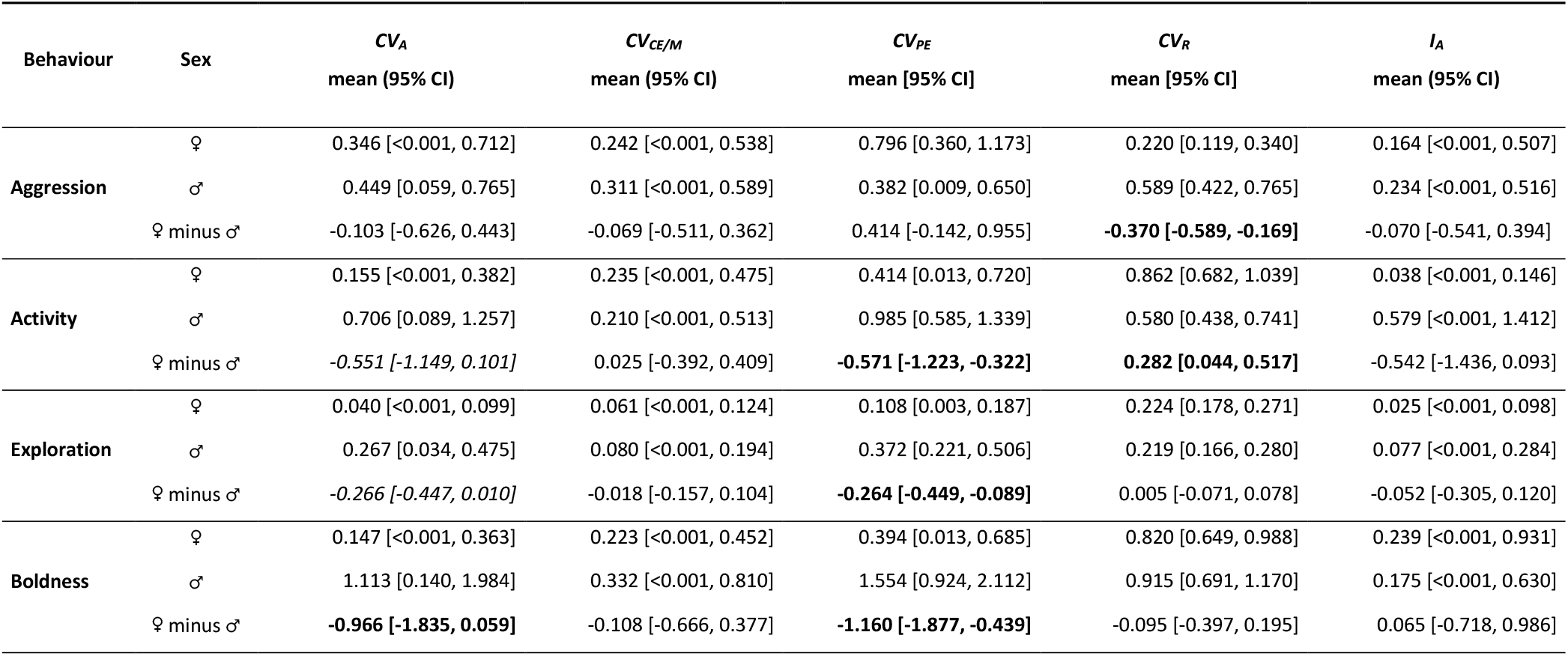
Estimates (posterior mean (95% credible interval) of the coefficient of additive genetic variance (*CV_A_*), common environment/ maternal effects variance (*CV_CE/M_*), permanent environment variance (*CV_PE_*), residual variance (*CV_R_*), and evolvability (*I_A_*). We used the raw behavioural data for the analyses.

| Behaviour | Sex | $CV_A$<br>mean (95% CI) | $CV_{CE/M}$<br>mean (95% CI) | $CV_{PE}$<br>mean [95% CI] | $CV_R$<br>mean [95% CI] | $I_A$<br>mean (95% CI) |
| --- | --- | --- | --- | --- | --- | --- |
| Aggression | ♀ | 0.346 [<0.001, 0.712] | 0.242 [<0.001, 0.538] | 0.796 [0.360, 1.173] | 0.220 [0.119, 0.340] | 0.164 [<0.001, 0.507] |
|  | ♂ | 0.449 [0.059, 0.765] | 0.311 [<0.001, 0.589] | 0.382 [0.009, 0.650] | 0.589 [0.422, 0.765] | 0.234 [<0.001, 0.516] |
|  | ♀ minus ♂ | -0.103 [-0.626, 0.443] | -0.069 [-0.511, 0.362] | 0.414 [-0.142, 0.955] | **-0.370 [-0.589, -0.169]** | -0.070 [-0.541, 0.394] |
| Activity | ♀ | 0.155 [<0.001, 0.382] | 0.235 [<0.001, 0.475] | 0.414 [0.013, 0.720] | 0.862 [0.682, 1.039] | 0.038 [<0.001, 0.146] |
|  | ♂ | 0.706 [0.089, 1.257] | 0.210 [<0.001, 0.513] | 0.985 [0.585, 1.339] | 0.580 [0.438, 0.741] | 0.579 [<0.001, 1.412] |
|  | ♀ minus ♂ | *-0.551 [-1.149, 0.101]* | 0.025 [-0.392, 0.409] | **-0.571 [-1.223, -0.322]** | **0.282 [0.044, 0.517]** | -0.542 [-1.436, 0.093] |
| Exploration | ♀ | 0.040 [<0.001, 0.099] | 0.061 [<0.001, 0.124] | 0.108 [0.003, 0.187] | 0.224 [0.178, 0.271] | 0.025 [<0.001, 0.098] |
|  | ♂ | 0.267 [0.034, 0.475] | 0.080 [<0.001, 0.194] | 0.372 [0.221, 0.506] | 0.219 [0.166, 0.280] | 0.077 [<0.001, 0.284] |
|  | ♀ minus ♂ | *-0.266 [-0.447, 0.010]* | -0.018 [-0.157, 0.104] | **-0.264 [-0.449, -0.089]** | 0.005 [-0.071, 0.078] | -0.052 [-0.305, 0.120] |
| Boldness | ♀ | 0.147 [<0.001, 0.363] | 0.223 [<0.001, 0.452] | 0.394 [0.013, 0.685] | 0.820 [0.649, 0.988] | 0.239 [<0.001, 0.931] |
|  | ♂ | 1.113 [0.140, 1.984] | 0.332 [<0.001, 0.810] | 1.554 [0.924, 2.112] | 0.915 [0.691, 1.170] | 0.175 [<0.001, 0.630] |
|  | ♀ minus ♂ | **-0.966 [-1.835, 0.059]** | -0.108 [-0.666, 0.377] | **-1.160 [-1.877, -0.439]** | -0.095 [-0.397, 0.195] | 0.065 [-0.718, 0.986] |

**Figure 2.**
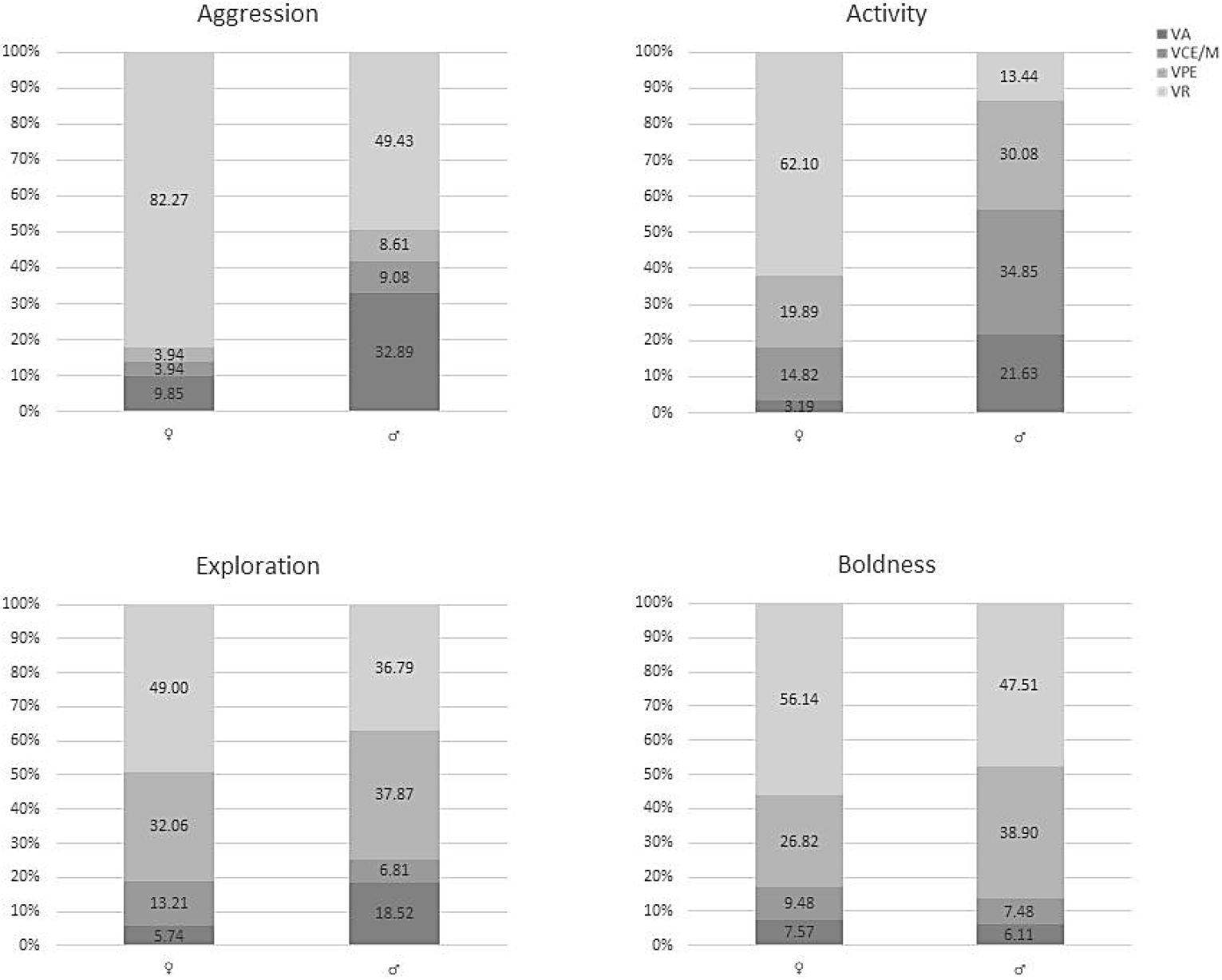
Variance components of behaviours – aggression, activity, exploration and boldness – in females and males. Stacked bars represent the total phenotypic variance (100%) decomposed into additive genetic variance (*V_A_*), common environment/ maternal effects variance (*V_CE/M_*), permanent environment variance (*V_PE_*), residual variance (*V_R_*). The numbers indicate the percentages of total variance explained by *V_A_*, *V_CE/M_, V_PE_, V_R_* in females and males.

### Activity

Males exhibit higher levels of expressed activity in the novel environment than females, however a sex difference is not significant (*female minus male*: post. mean difference = −0.250, 95% credible interval (CI) = [−0.543, 0.037,], p = 0.089; Figure 1). The quantitative genetic estimates, *V_A_*, *V_R_*, *V_CE/M_* and *V_PE_*, and *h^2^* for females and males are given in Table 1. Figure 2B shows how the *V_P_* in activity is partitioned into in each sex. The results suggest higher *V_CE/M_* and *V_PE_* in males compared with females (*V_CE/M_*, post. mean difference = −0.738, 95% CI [−1.323, −0.130]; *V_PE_*, post. mean difference = −0.599, 95% CI [−1.087, −0.092]). We also estimated higher *CV_PE_* in males (post. mean difference = −0.571, 95% CI [−1.223, −0.322]), whereas *CV_CE/M_* does not differ between the sexes (Table 2). Furthermore, males also have higher estimate of *V_A_* and heritability in activity than females, but sex differences are not statistically significant (*V_A_*, post. mean difference = −0.491, 95% CI [−1.185, 0.036]; *h^2^*, post. mean difference = −0.254, 95% CI [−0.627, 0.063]) though we note again that the CIs inform that we cannot rule out a true difference between these statistics. Similar results were obtained for *CV_A_* and *I_A_* (*CV_A_*, post. mean difference = −0.551, 95% CI [−1.149, 0.101]; *I_A_*, post. mean difference = −0.542, 95% CI [−1.436, 0.093]).

The posterior mean additive genetic covariance across sexes is estimated to < −0.001 with 95% CI between −0.109 and 0.103. The cross-sex genetic correlation is 0.012, 95% CI [−0.886, 0.990]. While Heidelberger and Welch’s diagnostic for the estimation of the additive genetic covariance across sexes passed Stationary test, it did not pass Halfwidth Mean test even with increased iterations, suggesting that estimation of the cross-sex correlation in activity is not reliable.

### Exploration

There are no sex differences in the expressed level of exploratory behavior (post. mean difference = – 0.047, 95% CI [−0.325, 0.226]; Figure 1). None of the quantitative genetic estimates; *V_A_*, *V_R_*, *V_CE/M_* and *V_PE_* differ significantly between the sexes (Table 1; Figure 2C). However, we found that males exhibit significantly higher *CV_PE_* compared to females (post. mean difference = – 0.264, 95% CI [−0.449, −0.089]). Furthermore, *CV_A_* is lower in females; the difference is not statistically significant, but based on the CI this difference cannot be ruled out (post. mean difference = −0.266, 95% CI [−0.447, 0.010]. Heritability estimates are 0.056, 95% CI [<0.001, 0.208] and 0.181, 95% CI [<0.001, 0.467] for females and males, respectively. We found no sex difference in either *h^2^* or *I_A_* (Tables 1–2). The estimate of the posterior mean additive genetic covariance across sexes is −0.003 with 95% CI between −0.115 and 0.113. The cross-sex genetic correlation is −0.013, 95% CI (−0.884, 0.847].

### Boldness

We found no sex differences in boldness levels (post. mean difference = – 0.106, 95% CI [−0.421, 0.196]; Figure 1). Females have higher *V_R_* than males (post. mean difference = 0.383, 95% CI [0.030, 0.734]), however, the two sexes do not significantly differ in *V_A_*, *V_CE/M_, V_PE_* and *h^2^* (Table 1; Figure 2D). Heritability estimates are 0.071, 95% CI [<0.001, 0.254] and 0.060, 95% CI [<0.001, 0.211] for females and males, respectively. We found however higher *CV_A_* and *CV_PE_* in male than in females (*CV_A_*, post. mean difference = −0.966, 95% CI [−1.835, 0.059]; *CV_PE_*, post. mean difference = −1.160, 95% CI [−1.877, −0.439]). The sex difference in *CV_A_* is not significant, but based on CI it can not be ruled out. The estimates of sex difference in *CV_CE/M_, CV_R_* and *I_A_* are not significant (Table 2). The posterior mean additive genetic covariance across sexes is 0.007, 95% CI [−0.051, 0.079]. The cross-sex genetic correlation is 0.082 with 95% CI between −0.731 and 0.871.

## DISCUSSION

Our quantitative genetic approach in a sexually-size dimorphic spider, *Larinioides sclopetarius*, explored sex-specific heritability estimates and cross-sex genetic correlations for several important behaviors. We reported (i) low to moderate heritability of personality traits – aggression, activity, exploration and boldness; (ii) that males show higher heritability estimates and additive genetic variances in aggression and activity than females and, further, that large sex differences in these statistics for these two behaviours cannot be dismissed. We detected no sex differences in the amount of genetic and environmental variances of exploration and boldness. Finally, iii) all traits have a mean cross-sex genetic correlation (*r_mf_*) close to zero, though it should be noted that the calculated cross-sex genetic correlations have broad 95% credible intervals in all assessed behaviors. Taken together, and also considering that empirical tests of sexual conflict that are based on the cross-sex genetic correlation for fitness are conservative (Connallon and Matthews 2019), our results imply that individual variation in aggression and activity might stem from sex-specific selection acting on these traits, and that the genetic architecture of these behaviors is sex-specific, allowing their sex-independent evolution.

We predicted to find a sex specific genetic architecture underlying aggressive behaviour. Our results show that males have both a higher additive genetic variance and a higher residual variance of aggression compared to females. Males also tend to have a higher heritability estimate and higher variances due to common environment/maternal and permanent environmental effects. The higher additive genetic variance of aggression implies that this trait has a higher potential to respond to selection in males compared to females. It has been suggested that traits closely related to fitness including life-history traits and sexually selected traits may have higher additive genetic and non-genetic variability than other traits (Houle 1992; Price and Schluter 1991; Rowe and Houle 1996; Merila and Sheldon 1999). This is consistent with our findings, to the extent that male aggression and fitness components are related in *L. sclopetarius*, as found previously (Kralj-Fišer et al. 2013). Under laboratory conditions, aggressive males sire more offspring, i.e. are more fertile, than non-aggressive ones, while aggression is not related to fecundity in females (Kralj-Fišer et al. 2013). In other words, intra-sex aggression is likely to affect fitness components in males to a higher extent than in females. We need to be cautious, however, as additional studies in wild populations are needed to draw these conclusions with confidence.

High levels of genetic variance in fitness traits and sexually selected traits would be expected if these traits represent larger mutation targets (Houle et al. 1996). Rowe and Houle (1996) suggested that high genetic variation in sexually selected traits, which are often costly to express and condition dependent, reflects the underlying genetic variance in condition. In *L. sclopetarius*, male-male combats that ultimately determine access to mates are common, meaning that it is likely that sexual selection has shaped intra-sex aggression in males. In females, aggression levels towards same-sex conspecifics, which is used to defend the territory and the foraging patch, is much lower. In both sexes aggression is subjected to trade-offs; overt aggression is costly due to injuries and deaths (Kralj-Fišer and Schneider 2012). The spiders for this study were weight-matched when tested for aggression, which prevents analysis of the relationship between body condition and aggression. In most spiders, however, larger males readily win the fights (Hoefler 2007). If this was also the case in *L. sclopetarius*, a sex difference in the amount of additive genetic variability may also imply that aggression is condition-dependent and that has been subjected to sexual selection in males, but less so in females.

We predicted that males, which exhibit higher repeatability in activity and exploration (Kralj-Fišer and Schneider 2012), should have higher heritability in these two traits (Falconer and Mackay 1996). As predicted, males tend to have higher heritability and additive genetic variances in activity compared with females; however, we found no such differences in exploration (Table 1). Sex differences in genetic variances in activity suggest that selection pressures acting on this trait might differ between the sexes, and implies a higher potential for a response in this trait to selection in males than in females. The result is not surprising given the life-style differences between adult orb-web females and males. Females are rather passive sit-and-wait predators, whereas males wander around actively searching for mates but cease web-building and foraging. It would be interesting to extend this kind of tests to the juvenile stages, when both sexes behave as sit-and-wait predators.

We found no sizeable differences in the estimates of sex-specific genetic variances and heritability of boldness (Table 1). These results imply a similar potential for evolutionary responses to selection in boldness in both sexes. Biologically, we might expect that males and females have a comparable propensity for risk-taking behaviors (boldness) if they equally increase their access to resources or chances of survival. Along the same lines, selection on boldness may not differ between the sexes.

Across species, most of the few previous studies investigating sex-specific genetic variances in aggression have found higher amounts of genetic variance for male compared to female aggression (Table 3). Again, this suggests that sexual selection, specifically male – male competition, shapes the genetic architecture of aggression in a sex-specific way in a variety of species. While the results on sex-specific genetic variances of activity and exploration are mixed, most studies found no sex differences in the amount of genetic variance of boldness (Table 3). Notably, if a sex difference in a behavior was detected, it consistently indicated higher additive genetic variances in males than in females (Table 3). This is in agreement with the findings of Wyman and Rowe (2014), who showed male-biased coefficients of additive genetic variance when analyzing sex differences in the amount of genetic variation across a wide range of traits and species. Additional studies are needed to pinpoint general patterns regarding sex-specific genetic effects in behavioral traits.

**Table 3.** Sex differences in additive genetic variance (*V_A_*), and heritability (*h^2^*) for behaviors related to personality.

| | | $V_A$ | $h^2$ | Reference |
| --- | --- | --- | --- | --- |
| Aggression | *Gryllus bimaculatus* | ♂ > ♀ | ♂ > ♀ | Han and Dingemanse 2017 |
|  | *Nuctenea umbratica* | no sex diff. | ♂ > ♀ | Kralj-Fišer et al. 2019 |
|  | *Larionioides sclopetarius* | ♂ > ♀ | ♂ > ♀ | this study |
| Activity | *Nuctenea umbratica* | no sex diff. | no sex diff. | Kralj-Fišer et al. 2019 |
|  | *Larionioides sclopetarius* | ♂ > ♀ | ♂ > ♀ | this study |
| Exploration | *Parus major* | no sex diff. | no sex diff. | van Oers et al. 2005 |
|  | *Gryllus bimaculatus* | ♂ > ♀ | ♂ > ♀ | Han and Dingemanse 2017 |
|  | *Larionioides sclopetarius* | no sex diff. | no sex diff. | this study |
| Boldness/risk-taking | *Parus major* | no sex diff. | no sex diff. | van Oers et al. 2005 |
|  | *Poecilia reticulata* | no sex diff. | no sex diff. | White et al. 2019 |
|  | *Larionioides sclopetarius* | no sex diff. | no sex diff. | this study |

Permanent environmental effects influenced the expression of activity, exploration and boldness to a high degree; i.e. they explained 20 – 40% of the total phenotypic variances. Comparably, a quantitative genetic study found high permanent environmental effects (65 – 85%) of exploration in sticklebacks (Dingemanse et al. 2012), and 18% and 5% of variance in exploration and aggression, respectively, in crickets (Han and Dingemanse 2017). Similar to our results, permanent environmental effects had a higher effect on exploration than aggression also in crickets (Han and Dingemanse 2017). High repeatability estimates in activity, exploration and boldness of *Larinioides* (Kralj-Fišer and Schneider 2012) may thus be explained largely by permanent environmental effects; i.e. paternal effects, epigenetics, non-additive genetic effects, (e.g., dominance variance (Wilson et al. 2010) and environmental effects that have long-term effects on phenotypes (nutrition during development, etc.). Hence, consistent individual differences in activity, exploration and boldness can be explained by state-dependent positive feedback. Notably, activity and exploration covary with spider weight in the offspring generation (Kralj-Fišer and Schneider 2012) suggesting that a part of individual variability in these behaviors may be explained by differences in weight.

Maternal effects, i.e. the mother’s influence on offspring phenotype beyond direct gene transmission, may importantly affect phenotypic variance within a population. Recent meta-analysis by Moore et al. 2019 reports that maternal effects account for about 10% phenotypic variation within populations, which is half as much as do additive genetic effects. These effects are similar between invertebrate and vertebrate species (Moore et al. 2019). In our study, hatchlings from the same clutch were reared in the same environment until the second molt disallowing to separate maternal from common environment effects. Thus, we accounted for the resemblance among siblings from the same mother stemming from the same early environment by including mother identity/common environment as a random effect in the models. This explained ~ 5 – 15% of the variance in aggression, exploration or boldness. While the percentage of variance in activity for females is also in this range, more than 30% of total phenotypic variance in activity was explained by maternal / common environment effects in males. These effects on activity are significantly higher in males than in females. Additional research on maternal effects underlying behavioral traits, and specifically personality traits, is necessary to generally assess the importance of these indirect genetic influences underlying individual behavioral variation.

Our heritability estimates of behaviors using the animal model approach differ considerably from the published results from mid-parent mid-offspring regression (Kralj-Fišer and Schneider 2012). While the previous analyses revealed “significant” heritability only for aggression, and a tendency for heritability in boldness (Kralj-Fišer and Schneider 2012), the animal model used here found that the investigated traits have low to moderate heritability (Table 1). The discrepancy in the results obtained through mid-parent mid-offspring regression versus animal model is not unusual (Kruuk 2004; De Villemereuil et al. 2013). Comparing studies that used both methods, Kruuk (2004) found heritability and standard errors estimated from animal model analyses to be lower than those from the parent–offspring regression or full-sib analyses. Kruuk (2004) attributed these differences to other sources of variance not accounted for in the simpler techniques. However, Åkesson and collegues 2008 criticized that conclusion, as Kruuk (2004) evaluated not only two approaches, but also different datasets. A subsequent comparison of the animal model and parent-offspring regression methods on the same data set found no general patterns (De Villemereuil et al. 2013). The differences between the two approaches are therefore difficult to assess (De Villemereuil et al. 2013). Our result on behavioral heritability using the same data set with different statistical approaches (Kralj-Fišer and Schneider 2012 vs. this study) implies that the animal model may be a more precise approach to reveal the genetic underlying of the measured traits than parent-offspring regression. The latter does not partition the components of variance in a trait, such as maternal, common environmental, permanent environmental effects, which is why the behavioral heritability estimates were higher (Kralj-Fišer and Schneider 2012). Notably, the present study clearly shows that genetic underpinnings of behaviors may largely differ between sexes, and thus, it underscores the importance of taking sex differences into account in quantitative genetic studies.

The mean additive genetic covariances across sexes in the behavioral traits were estimated to be around zero, yet the calculated cross-sex genetic correlations have rather large 95% credible intervals. We therefore note that we interpret the cross-sex genetic correlations estimates with caution, also due to rather small sample size. In accordance to our predictions, our results inform that *r_mf_* for aggression is likely lower than unity (0.170, 95% CI [−0.557, 0.852]), which would suggest mostly resolved intra-locus sexual conflict of this trait. This would imply sex differentiation in the genetic basis for aggression allowing for the independent evolution of this trait across sexes (Bonduriansky and Chenoweth 2009; Cox and Calsbeek 2009). Similarly, a previous study found a *r_mf_* for aggression significantly lower than one in the cricket *Gryllus bimaculatus* (Han and Dingemanse 2017). Our analyses indicate that *r_mf_* for exploration and boldness may be close to zero, which implies sex-specific genetic architecture and thus little genetic constraint on the sex-independent evolution in exploration and boldness too. In comparison, *r_mf_* estimates were close to unity for exploration in the cricket *Gryllus bimaculatus* (Han and Dingemanse 2017) and for risk-taking behaviors in the guppy, *Poecilia reticulata* (White et al. 2019).

## Conclusions

We report low to moderate heritability of behavioural traits – aggression, activity, exploration and boldness. We detected higher additive genetic variances in aggression and activity in males compared to females, but no sex differences in quantitative genetic estimates for exploration and boldness. The estimated mean cross-sex genetic correlation in all traits were close to zero suggesting these traits could have flexibility for sex-independent evolution. Taken together, our results imply that individual variation in aggression and activity might stem from sex-specific selection acting on these traits, and that the genetic architecture of these behaviours is sex-specific, allowing their sex-independent evolution.

## COMPLIANCE WITH ETHICAL STANDARDS

Conflict of Interest: The authors declare that they have no conflict of interest.

## ACKNOWLEDGEMENTS

We thank Katie L. Laskowski for her comments on the manuscript. SKF was granted a Humboldt Fellowship, and was supported by the Slovenian Research Agency (grant P1-0236). FGG was supported by a grant from the Spanish Ministry of Economy (CGL2016-76173-P) co-funded by the European Regional Development Fund. KLL was supported by a grant from the Deutsche Forschungsgemeinschaft (LA 3778/1-1). MK acknowledges support from the Slovenian Research Agency (grants P1-0255, J1-9163).

## Notes

### Competing Interest Statement

The authors have declared no competing interest.

